# MINSTED fluorescence localization and nanoscopy

**DOI:** 10.1101/2020.10.31.363424

**Authors:** Michael Weber, Marcel Leutenegger, Stefan Stoldt, Stefan Jakobs, Tiberiu S. Mihaila, Alexey N. Butkevich, Stefan W. Hell

**Author notes:** These authors contributed equally (Michael Weber, Marcel Leutenegger).

## Abstract

We introduce MINSTED, a stimulated-emission-depletion (STED) based fluorescence localization and super-resolution microscopy concept providing spatial precision and resolution down to the molecular scale. In MINSTED, the intensity minimum of the STED donut, and hence the point of minimal STED, serves as a movable reference coordinate for fluorophore localization. As the STED rate, the background, and the required number of fluorescence detections are low compared to most other STED microscopy and localization methods, MINSTED entails substantially less fluorophore bleaching. In our implementation, 200-1000 detections per fluorophore provide a localization precision of 1-3 nm in standard deviation, which in conjunction with independent single fluorophore switching translates to a ~100-fold improvement of far-field microscopy resolution over the diffraction limit. The performance of MINSTED nanoscopy is demonstrated by imaging the distribution of Mic60 proteins in the mitochondrial inner membrane of human cells.

---

To resolve fluorophores that are far closer than the diffraction limit, all lens-based fluorescence nanoscopy methods have to make adjacent fluorophores discernible during registration and identify their coordinates with high precision. The elegance of stimulated-emission-depletion (STED) microscopy^1,2^ derives from the fact that both tasks are performed in one go by the donut-shaped STED beam. By confining their fluorescence ability to a subdiffraction-sized region around its central minimum, the STED donut beam both singles out the fluorophores that happen to be located in this region and establishes their position. The fluorescence ability and therefore the region defined by the STED donut are well described by the effective point-spread-function (E-PSF) of the STED-microscope^3^, a Gaussian of full-width-half-maximum (FWHM) 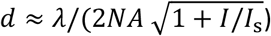. Here *λ*, *NA*, *I*, and *I*_s_ denote the wavelength of the STED beam, the numerical aperture of the lens, the focal peak intensity at the donut crest, and the intensity that reduces the fluorescence ability by half, respectively. Thus, scanning the sample with co-aligned, typically sub-nanosecond pulsed, excitation and STED beams separates fluorophores that are further apart than *d* and also locates them with standard deviation *σ*_*E*_ ~ 0.42*d*.

Interestingly, if *d* becomes as small as the fluorophore itself (1-2 nm), which is theoretically possible for *I* > 10^4^*I*_s_, all fluorophores will be prevented from fluorescing except the one that happens to be located right at the central donut minimum. At this conceptual limit without background, detecting just a single photon per fluorophore renders a perfect image, because a single detection within a given time span verifies the presence of a fluorophore at a coordinate perfectly defined by the donut. No other super-resolution fluorescence concept can make emitted photons as informative as STED microscopy and its close derivatives^4^.

Unfortunately, separating and locating emitters in one go comes at a cost. Since fluorescence blocking by STED typically entails intensities *I*_s_ ~ 1-10 MW/cm^2^, discerning fluorophores closer than *d* =20 nm requires *I* > 100*I*_s_ ~ 0.1-1 GW/cm^2^. Apart from the fact that applying such intensities to excited fluorophores promotes bleaching, donut minima are rarely < 0.01*I* in practice^3^. This means for *I* > 100 *I*_s_ that the intensity at the minimum exceeds *I*_s_, which leaves no fluorophore emitting and precludes fluorophore separation at distances well below 20 nm.

Here we introduce MINSTED nanoscopy, the first STED-based super-resolution fluorescence microscopy method that can provide molecule-size (1-3 nm) spatial resolution. This breakthrough has become possible by not requiring the STED donut to separate fluorophores (at small distances); its role is rather to establish the fluorophore’s position. While we give up some of the elegance of the original STED concept, we obtain the first fluorescence microscopy method whose resolution can be tuned from the diffraction limit down to the size of the fluorophores themselves. Compared to most other advanced STED and super-resolution methods^4^, MINSTED nanoscopy and the pertinent MINSTED localization entail less bleaching and reach the molecular scale with much fewer detected photons than popular camera based techniques.

To separate fluorophores at nanometre distances, MINSTED nanoscopy employs fluorophores that are transferred from an inactive (off) to an active (on) state and back. In the active state the fluorophore can be optically excited and de-excited by stimulated emission as in the concept called Protected STED^5^. However, in MINSTED nanoscopy only one fluorophore within a diffraction-limited region shall be switched on at any given time, meaning that its coordinate is initially unknown across diffraction length scales^6,7^. The subsequent localization with the STED beam is greatly facilitated by the fact that the central minimum of the donut defines a coordinate to which the unknown coordinate of the fluorophore can be related. In the following, we refer to the position of the donut minimum as the ‘donut position’. Since it can be steered with beam deflectors at sub-nanometre precision, the donut position can be used for finding the fluorophore position in a sample: the closer it is to the fluorophore, the lower is the STED probability and the more probable is fluorescent emission. Evidently, the donut position entailing minimal STED must be identical with the fluorophore coordinate; hence the name MINSTED.

In contrast to the related concept called MINFLUX^8^, searching for the donut position with minimal STED is tantamount to searching for the position where fluorescence is maximal. Yet this does not imply maximizing emission per se. Firstly, the absolute emission rate is freely adjustable via the excitation beam power. Secondly, and more importantly, placing the donut minimum on top of the fluorophore to maximize emission is neither required nor desired. Since the E-PSF is a Gaussian function, moving the E-PSF maximum in close proximity to the fluorophore does not provide the most precise localization per number of detected photons^8,9^. To find the peak of a Gaussian E-PSF it is in principle more photon-efficient to shift the peak aside and detect the rare and hence more position-informative photons generated at the Gaussian tail^8,9^. Unfortunately, the fluorescence photons from the tail are usually covered by background, rendering localization with diffraction-limited Gaussian excitation beams unattractive for most applications. In MINSTED, however, we narrow the E-PSF down, leave the diffraction-limit behind, and make all detected photons more informative in general. While a ‘STED microscopy of *d* = 1 nm’ is still hard to reach with normal fluorophores, the localization precision *σ* continues to scale with 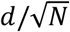. The number of detected photons *N* needed for reaching a certain *σ* decreases quadratically with decreasing *d*. Inserting the expression for *d* actually shows that 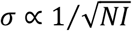. Thus, MINSTED can shift the demand for many photons from *N* to *I*, i.e. from the photon-poor fluorescence to the photon-rich donut beam. Only those fluorescence photons are required that indicate the position of the fluorophore with respect to that of the donut.

Importantly, whilst making the donut more intense and zooming-in on the fluorophore position, the donut can be translated so that the fluorophore always experiences intensities in the order of *I*_s_ and avoids the intensities *I* ≫ *I*_s_ found around the donut crest^10,11^. As we show in this paper, the unique combination of all these factors bestows MINSTED nanoscopy with molecule-size precision and resolution.

## Results

### MINSTED implementation, localization algorithm, and simulations

We implemented MINSTED in a confocal scanning microscope with electro-optic deflectors (EODs) and galvanometer mirrors for fast and slow scanning in the focal plane, respectively (Fig. 1a). After identifying an individual fluorophore by fleetingly scanning over the sample and estimating its position with 5-10 detections, the co-aligned excitation and STED beams were circled around a position estimate *C*_*i*_ with a radius *R*_*i*_ ≈ *d*_*i*_/2. Both *C*_*i*_ and *R*_*i*_ were updated after each photon detection *i* (Fig. 1b). Starting at *i* = 0 with a diffraction-limited E-PSF diameter *d*_0_ given for *I* = 0, the scan centre *C*_*i*_ was shifted by a fraction *α* of *R*_*i*_ toward the donut position, i.e. the E-PSF maximum, when detecting the next photon (*i* + 1). In other words, the donut minimum was moved tentatively closer to the fluorophore. This measure allowed us to sharpen the E-PSF by increasing the donut intensity *I*_*i*_ and reduce *R*_*i*_ by a factor *γ* at the same time, so that the ensuing smaller *d*_*i*_ left the ratio *R*_*i*_/*d*_*i*_ essentially unchanged. Therefore, despite the progressively higher *I*_*i*_ and steeper E-PSF slope, the fluorophore constantly experienced moderate donut intensities in the ballpark of *I*_s_ (Fig. 1c). A reduction of 3 % per photon detection (*γ* = 0.97) of *d*_*i*_ and a step size of 15% of the scan radius (*α* = 0.15) were typically used. We also set a limit *R*_*min*_ to the minimal radius and *I*_*max*_ to the highest donut intensity. During the circular scanning, the synchronously steered galvanometer mirrors ensured that the scan centre *C*_*i*_, i.e. the position estimate of the fluorophore, remained projected onto the confocal detector. As we zoomed-in on the fluorophore, the precision *σ* improved with decreasing *d*_*i*_, whilst the average emission rate remained largely constant.

**Fig. 1.**
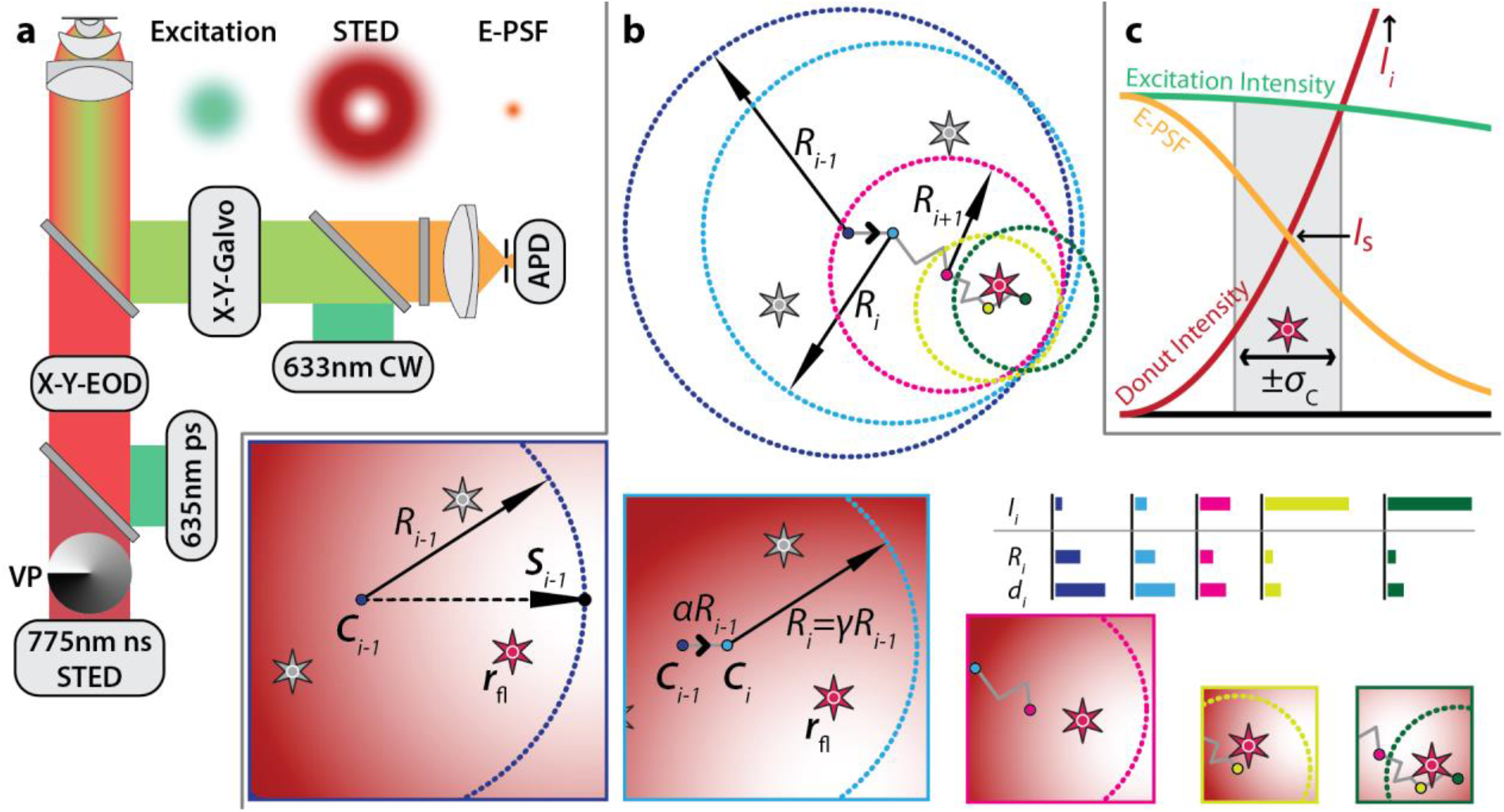
Principles of MINSTED localization. **a,** STED setup with co-aligned pulsed lasers for excitation and STED at 635 nm and 775 nm, respectively, and a vortex phase plate (VP) for helical phase modulation converting the STED beam into a donut; the inserts sketch the excitation and STED probability in the lens focal plane, along with that of fluorescence (E-PSF). The 633 nm CW laser was used for fluorophore pre-identification in the focal plane, while the X-Y-Galvo unit also maintained the optical conjugation of the confocal avalanche photodiode (APD) detector to the centre *C*_*i*_ of the circular scan performed by the electro-optical lateral deflector (X-Y-EOD). **b,** The active fluorophore (red among grey stars) was localized by circular X-Y-scans. For each photon detection *i*, the centre *C*_*i*_ was shifted by a fraction *α* of the radius *R*_*i*_ toward the donut minimum. Simultaneously, *R*_*i*_ and the FWHM *d*_*i*_ of the E-PSF were scaled by *γ* < 1. The centre *C*_*i*_ thus converges to the fluorophore position as indicated in the lower panels that also sketch relevant parts of the donut. Once a minimum radius *R*_*min*_ (yellow) is reached, only *C*_*i*_ is updated and the localization terminated after the fluorophore became inactive (*N* detections). The column diagrams illustrate the decrease of *R*_*i*_ and of *d*_*i*_ with increasing donut intensity *I*_*i*_. **c,** Normalized probability of excitation (green) and fluorescence detection (E-PSF, yellow) as a function of radial distance *ρ* from the focal point, along with a non-normalized intensity profile of the STED beam donut (red). Although *I*_*i*_ is constantly increased during the localization to sharpen the E-PSF, the intensity experienced by the fluorophore remains about *I*_s_ within the ±*σ*_C_ position range of the centre positions *C*_*i*_ highlighted in grey.

To assess the optimal ratio *R*_*i*_/*d*_*i*_, we simulated the precision *σ* expected for different *R*_*i*_/*d*_*i*_, intensity steps *I*_*i*_, and peak signal-to-background ratios (SBR) (Fig. 2). The SBR is a crucial parameter defined as the maximum detection rate from a fluorophore divided by a uniform detection rate in the sample. For zero background and a given number of detections *N*, a Gaussian E-PSF achieves higher precision when *R*_*i*_/*d*_*i*_ is large. However, in the practical range 5 < SBR < 50 and for *N* = 100 detections, *R*_*i*_/*d*_*i*_ = 0.5 is a better choice (Fig. 2a), because in the presence of background the value of *σ* becomes smaller when the emitter is closer to the E-PSF maximum and provides more photons. The step size *α* has several effects on the distribution of centre positions *C*_*i*_ and hence on the position estimate. Small *α* increase the *N* needed to reduce the distance between *C*_*i*_ and the fluorophore, and to converge to a final centre distribution (Fig. 2b). Larger *α* help approaching the fluorophore quickly, but the weaker correlation among the successive *C*_*i*_ bears the risk of not converging at low SBR. Furthermore, the reduction of *d*_*i*_ and *R*_*i*_ by a factor *γ* is tightly connected to the best step size *α*. Making *α* larger implies that 1 − *γ* can also be larger than 3%. Increasing *α* and decreasing *γ* entails that the number *N*_c_ of detections needed to reach the final *d*_min_ becomes smaller. For low SBR the risk of ultimately missing the fluorophore position grows, as expected (Fig. 2b; see also **Suppl. Fig. S1** to **Suppl. Fig. S4** and Suppl. Video V1).

**Fig. 2.**
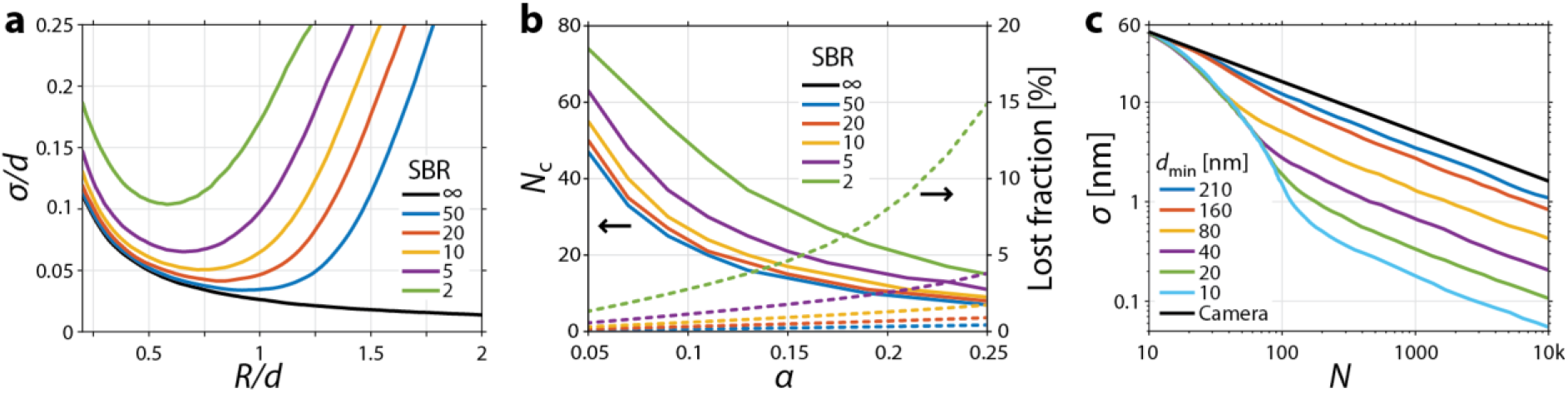
Simulation of MINSTED localization with *N* = 100 detected photons. **a**, Localization precision *σ* with different ratio of scan radius *R* to FWHM *d* of the STED microscope’s Gaussian E-PSF with SBR as parameter. While the hypothetical infinite SBR case calls for *R* maximization (black line), the presence of background enforces 0.5*d* ≤ *R* ≤ *d*. For large *R* the information provided by a single photon detection is masked by background, whereas for small *R* it is masked by the many other photon detections connected with an E-PSF maximum of finite *d*. **b**, Detections *N*_c_ necessary until the distribution of scan centre positions *C*_*i*_ (starting out with *d*_0_) converges to a final distribution (with static *d*_min_); percentage of simulations with centre positions *C*_*i*_ further than *d*_*i*_ away from the fluorophore and hence classified as lost. **c**, localization precision *σ* as a function of total number of detections *N* with *d*_min_ as parameter.

Altogether, a simulation of 500 localizations as a function of a finite number of detections *N* showed that the chosen parameters should provide robust and precise localization. As the donut scan centre homes in on the emitter, the *C*_*i*_ series serves as the permanently updated position estimate of the emitter. After reaching *d*_min_ at *i* = *N*_c_ and until reaching *i* = *N*, the coordinate average 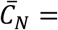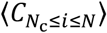 was our localization result. The simulation reveals the importance of the counts up to about *N*_c_ for zooming-in and reducing the *N* required for a certain *σ*. It also confirms that *σ* scales with 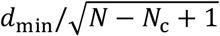 (Fig. 2c). Concretely, *d*_min_ = 40 nm is predicted to yield *σ* ≈ 3 nm with only *N* = 100 photon detections.

### Experimental MINSTED localization precision

To test these predictions, we localized immobilized individual ATTO 647N fluorophores on coverslips^12^ using MINSTED with *d*_min_ ≈ 40 nm. Driven by each detection *i*, the scan centre progressed toward the fluorophore and ultimately meandered around the estimated final coordinate (Fig. 3a,b and Suppl. Video V2). Recording many of these traces for many fluorophores allowed us to explore the attainable precision. The fluorophores were localized multiple times and the localization precision was analysed between the different localizations of the same molecule. To attribute localizations to individual fluorophores, we clustered localizations that were ≤ 25 nm apart. Only sets with more than five localizations were analysed and the scan centres *C*_*i*_ were regarded as the fluorophore coordinate estimates, as in the simulations. Once *d*_min_ = 40 nm was attained, the ‘meandering’ positions *C*_*i*_ were averaged to 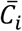 until the specified *N* and hence 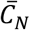 was reached. Within each localization cluster, the estimated final coordinates were calculated at multiple photon numbers to establish *σ* as a function of *N*.

**Fig. 3.**
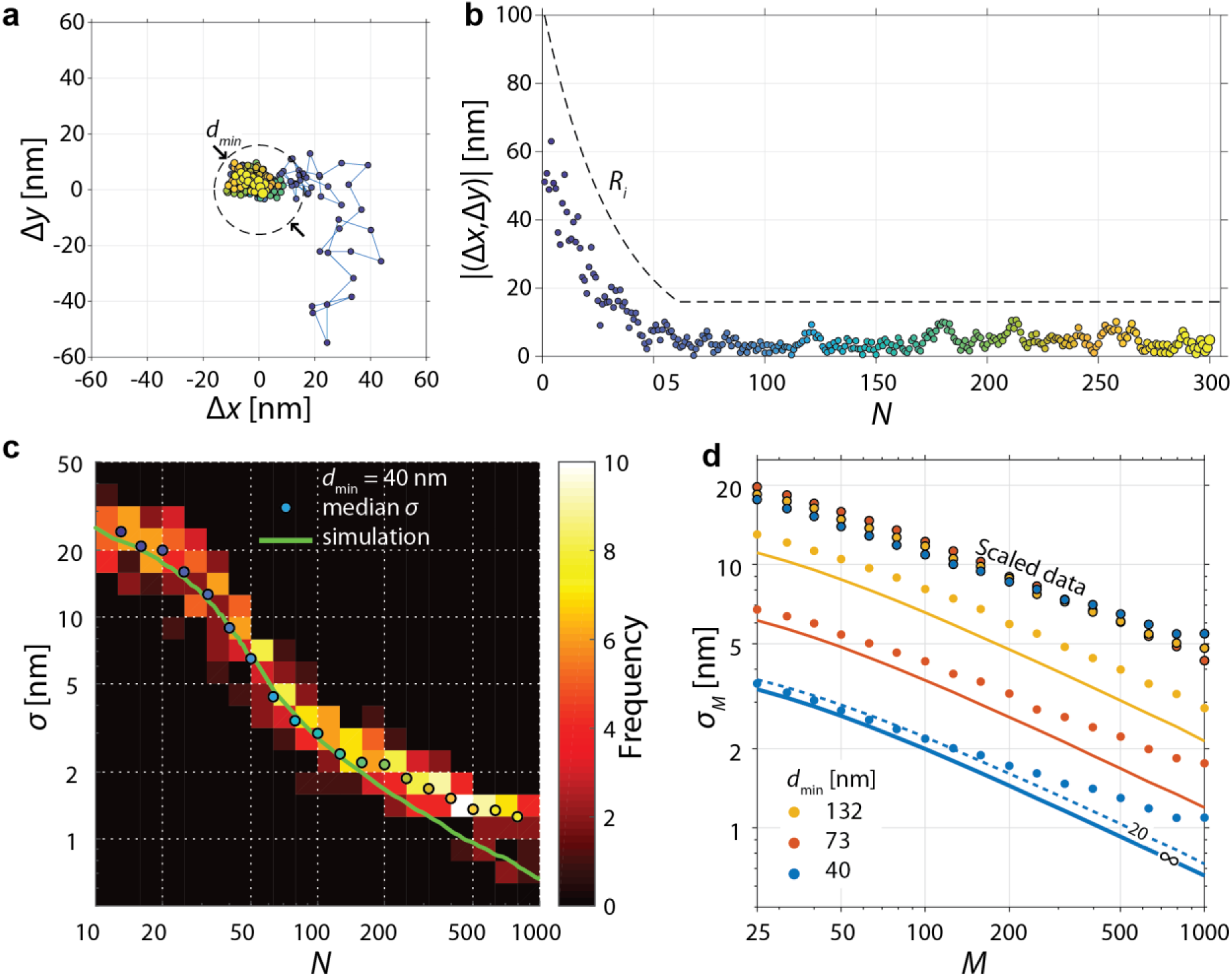
MINSTED localization of single fluorophores. **a**, Localization trace from first *i* = 1 (blue) to last detection *i* = 300 (yellow) with final scan circle (dashed line) around the estimated (*x*, *y*) position. **b**, Scan radius *R*_*i*_ (dashed line), distance (Δ*x*, Δ*y*) from the final estimated position to the scan centre *C*_*i*_ (points) from *i* = 1 (blue) to *i* = 300 (yellow). **c**, Histogram of precision *σ* of grouped localization traces and their median *σ* showing good agreement with simulation. **d**, Measured precision *σ*_*M*_ (derived from segments of *M* detections measured after *d*_min_ had been reached) showing how the increase of STED donut power improves the precision in linear proportion to *d*_min_, which is also confirmed by the overlap of data points when all points are scaled to *d*_min_ = 200 nm for comparison. Note the logarithmic display in **c** and **d**.

Our experiments show that *σ* decreases rapidly with decreasing *d*_*i*_ until *d*_min_ is achieved at *N*_c_ ≈ 60 (Fig. 3c). For *N* > *N*_c_, the precision *σ* follows the 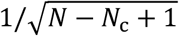 dependence (compare Fig. 2c) until it deviates from the simulation at about *σ* < 2 nm. This deviation is likely due to residual drifts of the fluorophore and/or the setup. The measured *σ* at around *N* = 10 is slightly better than the previously simulated values, because the 5-10 detections gained from the initial fluorophore identification by galvo-scanning provided *σ*_0_ ≈ 60 nm right at the outset. Consideration of this *σ*_0_ resulted in an excellent agreement between the simulated and experimental *σ* as a function of *N* (Fig. 3c). Since fluorophores moving substantially less than the standard deviation *σ*_C_ of *C*_*i*_ cannot be excluded, we can safely assert that in our experiments MINSTED reached *σ* = 2-3 nm with just *N* = 200 detections.

Next, we measured *σ* obtained after *d*_min_ had been reached. Since the total number of detections before bleaching typically exceeded 1000 per fluorophore, we split the resultant *C*_*i*_ traces into segments of different sizes *M* and calculated the standard deviations *σ*_*M*_ of the localization in these segments. To avoid boundary artefacts, we explored the range *N* − *N*_c_ > 25. In agreement with the simulations, the measurements again followed the 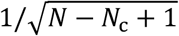 relation and the linear dependence on *d*_min_ (Fig. 3d and **Suppl. Fig. S5**). To highlight the latter, we also scaled the measured *σ*_*M*_ to *d*_min_ = 200 nm so that any difference from the linear dependence could be noticed in the overlay. At *σ* < 3 nm, the measured *σ* deviates from the simulations as before. Yet the data shows that at *d*_min_ = 40 nm, 1000 detected photons yield molecule-size precisions *σ* ≈ 1 nm. If residual movements of the stage or the fluorophore could be avoided, ~500 detections at SBR = 20 would suffice for *σ* ≤ 1 nm. Indeed, comparison of the measured precision with that simulated for the ideal SBR = ∞ case shows improved agreement for smaller *d*_min_, indicating that the STED donut not only improves the information of the detected photons by confining their origin in space, but also by suppressing background.

### MINSTED fluorescence nanoscopy in cells

Separation of emitters in MINSTED nanoscopy requires fluorophores that can be transferred from a lasting state that is non-responsive to excitation light into a semi-stable state leading to fluorescence upon excitation. Silicon rhodamine (SiR) fluorophores with two unsubstituted photoactivatable *ortho*-nitrobenzyl (ONB) caging groups proved suitable for MINSTED because photoactivation at 355 nm wavelength activated the SiR fluorophores enabling STED at 775 nm wavelength with no concurrent two-photon activation^13^.

To demonstrate its potential for biological imaging, we used MINSTED nanoscopy to image the mitochondrial inner membrane protein Mic60 in chemically fixed human cells^14^. The mitochondrial inner membrane folds into cristae, large membrane invaginations that increase the surface of this membrane. Mic60 is enriched at crista junctions^15^, which are round or slit-like structures that connect crista membranes with the mitochondrial inner boundary membrane that parallels the outer mitochondrial membrane.

Immunolabelling of cultured human U2OS cells with ONB-2SiR-labeled primary anti-Mic60 antibodies allowed us to compare MINSTED nanoscopy with confocal and STED images recorded after activation of ONB-2SiR (Fig. 4). Confocal microscopy was unable to provide details of the distribution of Mic60 in mitochondria (Fig. 4a). Featuring a resolution of about 60 nm, the recorded STED images demonstrated a clustering of Mic60, but failed to resolve individual proteins (Fig. 4b). In contrast, MINSTED accomplished this feat (Fig. 4c,d), recording 1.8–2.4 localizations per second and resolving individual fluorophores with a median precision of *σ* = 2.1 nm (Suppl. Fig. S6).

**Fig. 4.**
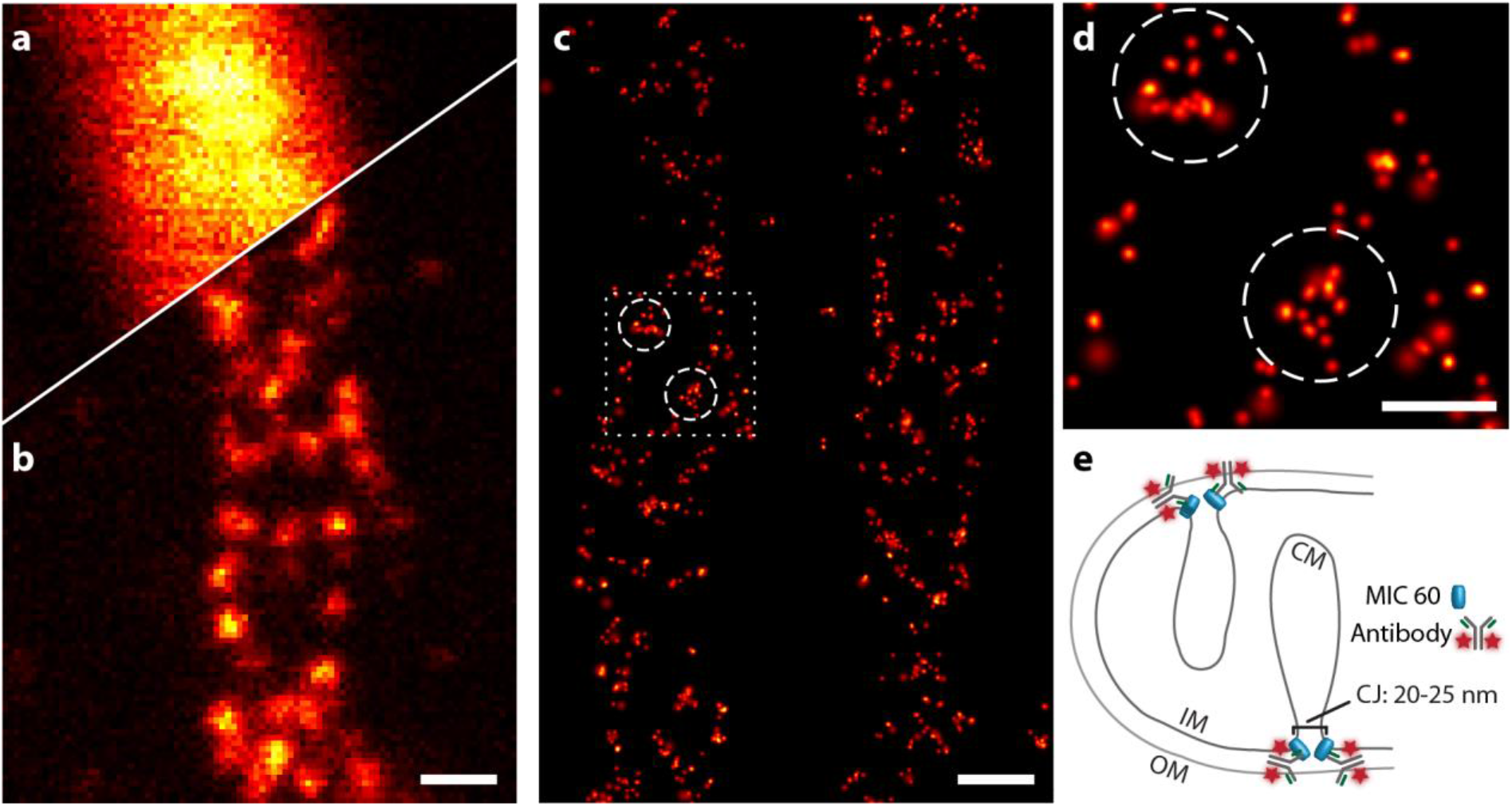
MINSTED nanoscopy of mitochondrial protein Mic60. **a,** Confocal and **b**, STED image with *d* ≈ 60 nm of the same mitochondrion taken after simultaneous activation of all fluorophores. **c,** MINSTED nanoscopy image of similar mitochondria resolving the Mic60 clusters (3607 localizations acquired in 33 minutes, 1463 localizations with *N* − *N*_c_ ≥ 250 detections, *d*_min_ = 54 nm). **d,** Excerpts of data as indicated in **c**. **e,** Cartoon of the presumed localization of Mic60 in the mitochondrial inner membrane. IM: inner membrane; OM: outer membrane; CM: crista membrane; CJ: crista junction. Scale bars: **a**–**c,** 200 nm; **d,** 100 nm.

For this study, we relied on primary antibodies that were labelled by azide modification of the glycans on the antibody heavy chain, so that the distance between the antibody binding site and the fluorophore was as small as 6-10 nm (PDB ID code: 1HZH^16^). The localization precision of individual fluorophores was three times higher than this distance, highlighting the limits set by the labels on extracting biological information at the single digit nanoscale. Since fluorescence microscopy cannot reveal anything but the fluorophores in the sample, our results show that MINSTED reaches the conceptual limits of this imaging modality. Nevertheless, our two-dimensional MINSTED data provides valuable insights about the nanoscale distribution of Mic60 in mitochondria. We repeatedly recorded a circular arrangement of Mic60 in mitochondria using MINSTED (Fig. 4d), which is in excellent agreement with the current understanding of Mic60 forming small ring-like assemblies at cristae junctions^17^. With future three-dimensional implementations of MINSTED, complex structures such as those found in the mitochondrial inner membrane should be accessible in detail.

## Discussion and Conclusion

Under the provision that adjacent fluorophores are individually active, MINSTED nanoscopy can deliver molecule-scale resolution like its MINFLUX counterpart. However, in MINSTED a resolution of *σ* = 2 nm or *d* = 4.7 nm is attained with a total of just 200 detections in close agreement with the simulations. The reason is that the STED donut suppresses spurious signal from the neighbourhood of the targeted fluorophore, rendering the MINSTED images (Fig. 4) almost background-free.

Another strength of the described MINSTED implementation is that the photon-by-photon update of the localization removes virtually all bias due to inaccurate assumptions on background or donut shape. Furthermore, the unequivocal repositioning of the donut centre in the right direction allows for an aggressive reduction of *d*_*i*_ and hence also of *N*. Once *d*_min_ is reached, each subsequent photon refines the scan centre *C*_*i*_ and lowers the uncertainty on the position estimate 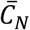. In fact, continuous updating of *C*_*i*_ tracks the fluorophore until it bleaches or switches off. As it provides the most photon-efficient localization to date, MINSTED will be useful for tracking rapidly moving emitters. Our MINSTED protocol can be further refined by dynamically adapting *R*_*i*_/*d*_*i*_ in response to the background. Evidently, future MINSTED research will include multiple colour channels using spectrally shifted fluorophores, 3D recordings using 3D donuts, and technically more sophisticated implementations with adaptable donut arrays or sets of standing waves.

The selective spatial targeting of the donut minimum constitutes a fundamental difference from earlier applications of STED microscopy to single fluorophore localization, whereby the donut is scanned laterally across the focal plane to map out the E-PSF centroid rendered by each fluorophore^12,19,20^. This established combination of single fluorophores and STED works reliably only for bleaching-resilient emitters, such as NV centres^19^, or for low donut intensities, because the intense donut crest typically bleaches the fluorophore before the whole E-PSF is acquired. Moreover, precise rendering of the E-PSF is typically compromised by the fluorophores’ tendency to blink. In MINSTED, although the donut intensity is constantly increased, bleaching and blinking aggravation is avoided. For attaining nanometre spatial resolution, MINSTED nanoscopy requires neither intensities > 10^4^*I*_s_ nor donut minima < 1 % since the on/off separation of spatially tight fluorophores is not performed by the donut but the on/off switching of individual fluorophores. Yet, the donut brings about the advantage that it additionally assists the on/off separation at subdiffraction length scales.

Seeking a fluorescence minimum, as in MINFLUX, has a conceptual advantage over localizing with a Gaussian E-PSF unless background comes into play. When narrowing the search range by increasing the donut intensity, the excitation donut in MINFLUX^8^ is more prone to worsening background levels than is the STED donut in MINSTED. In fact, we found that higher STED donut intensities at 775 nm keep the background low even for small *d*_min_. For this reason, MINSTED is currently on par with or even outperforms MINFLUX in key aspects.

In most of our MINSTED imaging, *d*_min_ was not reduced below 40 nm because a higher STED beam power would have increasingly destabilized the system by heating. Requiring a STED beam is an added complexity of MINSTED as compared to MINFLUX, but the precision *σ* achievable with either of the two molecule-scale resolution approaches will ultimately depend on the background. In any case, by featuring excellent background suppression, MINSTED should become substantially faster and handle higher densities of fluorophores than most super-resolution methods in the future.

Finally, the introduction of MINSTED underscores that the idea of optically injecting a movable reference coordinate is transformative in the art of localization of emitters. In conjunction with on-off state separation, MINSTED enlarges the scope of far-field fluorescence nanoscopy with molecule-size resolution, which due to its 100-fold improvement over the diffraction limit is poised to break new grounds.

## Methods

Methods and any associated references are available in the online version of the paper.

## Acknowledgments

We thank Roman Schmidt (now Abberior Instruments GmbH) for contributions to early versions of the software and the setup. We are also grateful to Frank Werner, Thomas Staudt and Jan Keller-Findeisen for discussions on the localization statistics and for complementary calculations, as well as to Klaus Gwosch and Francisco Balzarotti for discussions about MINFLUX. Taukeer Khan, Heydar Shojaei and Vladimir N. Belov supported us with the design and synthesis of fluorophores. We acknowledge Ellen Rothermel for preparing samples, Tanja Gilat and Marco Roose for technical support. Funding by the German Federal Ministry of Education and Research (BMBF) in the project “New fluorescence labels for protected- and multi-color-STED microscopy (STEDlabel)” (no.13N14122, to SWH) and by the European Research Council Advanced Grant 835102 (to SJ) is gratefully acknowledged. TSM was supported by a Fulbright Research scholarship.

## Author contributions

ML and MW designed and implemented the specific localization algorithm and performed the simulation analysis with critical input from SWH. ML and MW built the setup and ML wrote the software, including the real-time control of the setup. MW prepared the samples and performed the measurements. ANB synthesized dyes for preliminary tests. TSM explored various labelling techniques. MW and ML analysed the data with feedback from SWH. SWH outlined fundamentals of the MINSTED concept, initiated and supervised its exploration. SWH, ML and MW wrote the manuscript. All authors contributed to the manuscript and the supplementary information either through discussions or directly.

## Competing financial interests

SWH benefits from intellectual property on the described localization and nanoscopy owned by the Max Planck Society.

## Methods

### MINSTED setup

The setup consisted of an epi-fluorescence microscope with a dual-channel confocal laser scanning system using a Leica 100×/1.4NA oil-immersion objective lens. Two galvanometer mirrors and a pupil relay optics allowed for rapid beam scanning over a quadratic sample area of about 100 μm extent (x,y). A continuous-wave (cw) HeNe laser provided fluorescence excitation at 633 nm wavelength for rapid overview. A single-photon counting module detected the fluorescence light in the 650–750 nm range. A confocal pinhole of 0.5 Airy units diameter blocked out-of-focus light. For STED microscopy and single-molecule localization, an additional illumination path without moving parts was implemented. Two electro-optic deflectors with pupil relay systems featured beam scanning within a square image area of about 2.6 μm extent. A 635 nm pulsed diode laser delivered excitation pulses of about 100 ps duration, whereas a 775 nm pulsed fibre laser provided STED pulses of about 1 ns duration. A vortex phase plate imprinted a 2π phase ramp on the phase front of the STED beam and a polarization controller converted it to circular polarization in order to shape the STED beam into a donut profile. A diode laser at 355 nm wavelength illuminated the STED image area to photoactivate fluorophores. All laser beam powers were modulated with short response times of several μs. The sample was mounted on an X-Y-Z-piezo positioning stage whose position was locked by a sample tracking system. For this purpose, the position of fiducial markers was monitored with infrared light from a super-luminescent light emitting diode and fast CMOS cameras. The tracking system issued the closed-loop control signals to cancel sample drift. The MINSTED microscope was fully controlled by an FPGA board and a custom control program. Our software ran diffraction-limited overview scans using only the galvanometer beam scanner as well as high-resolution STED image scans and single-molecule localizations using both scanners synchronously. A graphical user interface allowed defining the measurement parameters and retrieving the measurement results.

### Immobilization of Atto 647N fluorophores

Atto 647N molecule were sparsely distributed and immobilized on cover slides as described in ref. 8. A flow channel, consisting of a cleaned coverslip glued to a microscope slide with double-sided scotch tape, was rinsed with 100 μl phosphate-buffered saline (PBS, 137 mM NaCl, 2.7 mM KCl, pH 7.4). The channel was filled with 15 μl biotinylated BSA (biotinylated bovine serum albumin, A8549, Sigma Aldrich, St. Louis, MO, USA) 0.5 mg/ml in PBS. After 4 min incubation, the channel was flushed with 100 μl PBS and filled with 15 μl Streptavidin (11721666001, Sigma Aldrich, St. Louis, MO, USA) 0.5 mg/ml in PBS. After an incubation time of 4 min, the channel was flushed with 100 μl PBS and filled with 15 μl 200 pM hybridized Biotin-DNA/Atto647N-DNA in PBS^8^. After 4 min incubation, the channel was flushed with 100 μl PBS and filled with 0.01 % (w/v) Polylysin (P8920, Sigma-Aldrich, St. Louis, MO, USA) in PBS for 10 min. After flushing with 100 μl PBS, the channel was filled with 15 μl freshly diluted silica shelled silver nanoplates (SPSH1050, nanoComposix, San Diego, CA, USA) 2.5 μg/ml in PBS. After 10 min incubation the channel was flushed with PBS again, filled with 15 μl ROXS buffer^21^ and sealed with epoxy glue (Hysol, Locktite).

### Antibody conjugation

The labelling of antibody using glycan modification and strain promoted click chemistry and the synthesis of the used dye was described previously^13^. In short, the rabbit monoclonal antibody (ab245764, Abcam, Cambridge, MA, USA) was modified with azide groups using a commercial enzyme system (GlyClick, Genovis, Lund, Schweden). After the modification, 250 μg antibody in 200 μl tris-bufferd saline (TBS, 20 mM Tris HCl, 150 mM NaCl, ph 7.6) was mixed with 50 μg DMF containing 50 μg DBCO-dye and stirred overnight. The free dye was removed using a phase extraction by adding 600 μl distilled water, 90 μl saturated (NH_4_)_2_SO_4_ solution and 900 μl tert-butanol, vortexing and separating the phases after a short centrifugation pulse. The aqueous phase (about 600 μl) was diluted using 600 μl TBS. The labelled antibodies were aliquoted and stored at −20°C.

### Cell labelling

The human Osteosarcoma cell line U-2 OS was obtained from the European Collection of Authenticated Cell Cultures (ECACC, Porton Down, Salisbury, UK; Cat no. 92022711, Lot. 17E015) and cultivated on coverslips in McCoy’s medium (Thermo Fisher Scientific, Waltham, MA, USA) supplemented with 10 % (v/v) fetal bovine serum (Thermo Fisher Scientific, Waltham, MA, USA), 1 % (v/v) sodium pyruvate (Sigma Aldrich, St. Louis, MO, USA) and Penicillin-Streptomycin (Sigma Aldrich, St. Louis, MO, USA). The cells were fixed using 8 % (w/v) Paraformaldehyde in PBS for 5 min, permeabilized with 0.5 % (w/v) Triton X-100 for 5 min and quenched with 100 mM NH_4_Cl in PBS for 5 min. The fixed cells were washed with PBS, blocked with 2 % (w/v) bovine serum albumin (BSA) in PBS and treated with the primary antibody in the same buffer for 1 h, washed with 2 % (w/v) BSA in PBS, treated with a secondary goat anti-rabbit antibody conjugated with Alexa 647 for MINSTED and washed with PBS.

### Cell imaging

The confocal and STED images were recorded on a commercial Abberior Instruments Expert Line microscope equipped with a 775 nm 40 MHz STED and a 640 nm excitation laser after activation with a spectrally broad 405 nm LED as described in reference ^13^. For MINSTED, the labelled cells were incubated with freshly diluted silica shelled silver nanoplates in PBS for 20 min and then washed with PBS. The samples were mounted with buffer (20 mM Hepes, 150 mM NaCl, pH 7) using TwinSeal (Picodent, Wipperfürth, Germany). Prior to MINSTED, the cells were selected based on the counterstain signal and the Alexa 647 dyes were bleached using low power STED light. The localization routine was started without excitation laser to equilibrate the temperature in the immersion oil and sample, which were warmed up by the STED laser. After 10 seconds, the excitation laser was enabled and the caged dyes were sparsely activated using 355 nm light when searching for another active fluorophore. Over the duration of the measurement, the UV laser power was slowly increased to keep the activation rate constant. The imaging was stopped when no further molecules could be activated.

### Data Analysis

The localizations were analysed based on the centre positions 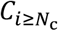 at *d*_min_. The localizations were further selected with a maximum filter on the standard deviation *σ*_C_ of the *C*_*i*_, together with a minimum filter on the number of detected photons *N*. The precision of each localization was estimated as described in the supporting information and validated by simulations (**Suppl. Fig. S3**). The image was rendered with the estimated precision lower-bounded to 3 nm.

## Supplementary information

### Supplementary text

#### MINSTED localization method

Image scans were performed until the total number of detected photons of the current and previous columns and rows exceeded a threshold *n*_ON_ of typically 5-10. When the threshold was exceeded, the image scan was interrupted and the detections at the last image positions were used for an initial estimation 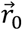 of the fluorophore position 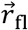 by a weighted average. A circular scan was started, whose initial centre position 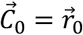 and whose radius *R*_0_ ≈ *d*_0_/2 was approximately half the PSF diameter as illustrated in Fig. 1b. During the localization, the centre converged towards the fluorophore position. Thus, the fluorophore was exposed to a moderate STED intensity because it was kept within a distance *d* from the donut position 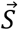 most of the time, typically at about the scan radius *R* (Fig. 1c). Upon detection of photon *i* at the donut position 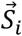, the centre, scan radius and E-PSF diameter were updated immediately. The centre was moved towards 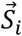 by a fraction *α* and the scan radius and E-PSF diameter were scaled by *γ* < 1. The scanning continued around the new centre 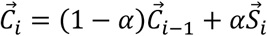 with radius *R*_*i*_ = *γR*_*i*−1_ and d*i* = *γ* d_*i*−1_ until the next photon was detected and the update repeated. The scan radius and the E-PSF diameter were decreased only until reaching their preset lower limits, whereas the centre position was updated throughout. The localization was terminated if less than a minimum of *n*_OFF_ photons were detected within a time interval *τ*_OFF_.

The real-time FPGA control logic was kept simple and lean. All localization traces consisting of centre positions, donut positions, detected photons and detection times were transferred to the computer for storage and further evaluation using MATLAB and custom analysis tools. The fluorophore position after *N* photon detections was estimated by the last centre position for *N* ≤ *N*_c_, where *N*_c_ is the number of detections to reach the smallest scan radius and sharpest E-PSF. For *N* > *N*_c_ the fluorophore position was estimated by the average centre positions during the remaining photon detections.

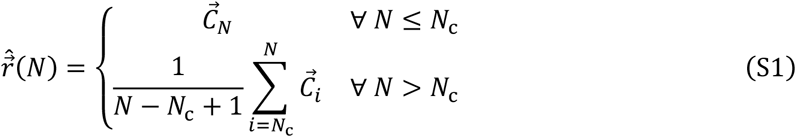

#### MINSTED localization uncertainty

Due to the continuous update of the centre position, the localization uncertainty – this means the Cramer-Rao bound (CRB) – could not be easily obtained analytically for the interesting case of many photon detections. Instead, we repeatedly simulated the localization of a fluorophore and estimated the localization uncertainty by the root mean squared (RMS) localization error versus the number of detected photons *N*.

Given the estimates 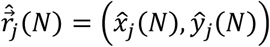 of the fluorophore position after *N* detections for *j* = 1 … *K* runs of the simulation, the localization uncertainty along *x* and *y* was estimated by the root mean square errors along the coordinate axes:

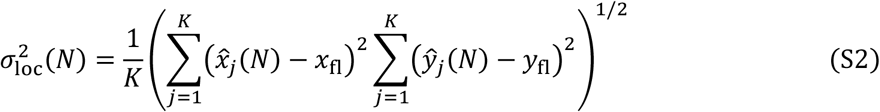

Simulated localizations that ended prematurely after *i* < *N* detections or whose positions were off by more than five times the median error of the simulations were flagged as failures for the remaining detections. Equation (S2) was evaluated for the successful localizations only. Fig. 2a–c, Fig. 3c,d illustrate localization uncertainties with simplified fast calculations, whereas **Suppl. Fig. S1** and **Suppl. Fig. S2** illustrate results based on complete simulations including the detection times and the termination of the fluorophore’s active state.

For the experimental localizations, the true fluorophore position and thus the accuracy is unknown. Based on simulated localizations, we found that the estimation of the localization precision similar to camera-based localizations is unreliable due to the history of the centre trace, which introduces a varying degree of correlation among the centre positions. Instead, we estimated the precision of an individual localization by sub-sampling. Therefore, we split the centre trace in groups of different sizes *M*, estimated the fluorophore position for each of these groups, and extrapolated the precision from the groups to the entire localization.

Given *N* ≫ *N*_c_ detected photons, we calculated for consecutive groups of *M* = [10^{1.5,1.6,1.7,…^ ^}^] ≤ (*N* − *N*_c_ + 1)/5 detections the standard deviations *σ*_*M*_(*M*) of the groups’ mean centre positions. We then estimated the precision of the localization by extrapolating the relation

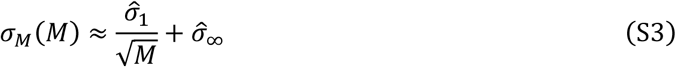

to *M* = *N* − *N*_c_ + 1. Here, 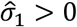 is approximately the standard deviation among uncorrelated centre positions and 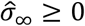 is an empirical offset, both obtained by least-squares decomposition of *σ*_*M*_(*M*).

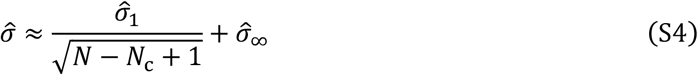

**Suppl. Fig. S3** compares the localization uncertainty obtained from numerous simulated localizations with the estimated precision extracted from the individual localizations. The extracted precision provides a reasonable and rather conservative estimate of the true uncertainty.

#### Simulated localization

The fluorophore was placed at 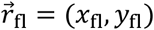 and the localization started with a centre position 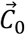 that was normally distributed in the (*x*, *y*) plane around this position with a standard deviation 90 nm. Hence, the initial centre distribution approximated the profile of the confocal fluorophore image used for searching the fluorophores.

A 2D Gaussian PSF with peak detection rate *ɛ* and a background detection rate*β* were assumed; that is a peak signal-to-background ratio SBR = *ɛ*/*β*. The average detection rate 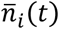 at donut position 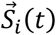 was therefore

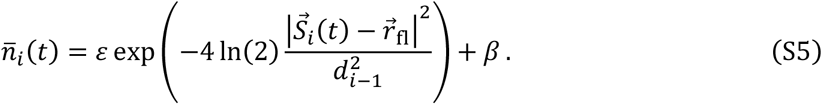

The scan trajectory leading to the *i*^th^ detection was determined by the centre position 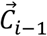 and the radius *R*_*i*−1_ after the previous detection.

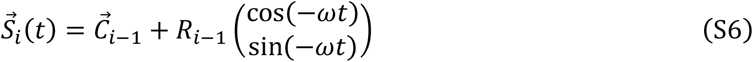

For each photon detection *i* = 1 … *N* an exponentially distributed number *m*_*i*_ with average value 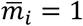 was drawn. The detection time *t*_*i*_ was then determined by integrating the signal 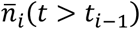 along the scan trajectory 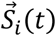 until its cumulative value reached *m*_*i*_.

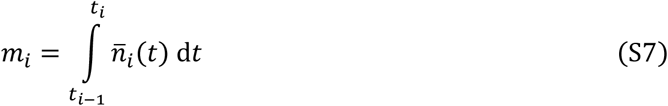

Upon each detection, the E-PSF was sharpened until reaching the minimal value d_*min*_ found for the maximal STED beam power. The scan radius was reduced equally until reaching its minimum *R*_*min*_.

**Suppl. Fig. S4** illustrates that the donut position 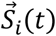 stays from the fluorophore at a narrow distance distribution centred on the scan radius during the entire localization. Therefore, the fluorophore is exposed only to a moderate STED intensity, which lowers photobleaching by the donut as compared to conventional raster-scanned STED imaging.

#### Camera-based localization uncertainty

The theoretical localization precision for single-molecule localizations by analysing camera images is given by Thompson et al.^22^ and Mortensen et al.^23^ as

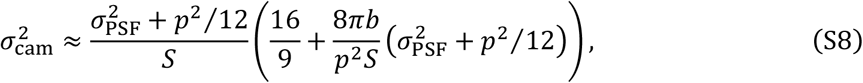

where *S* is the total number of detected signal photons; *p* is the pixel size; *b* is the average background per pixel without dark counts; *σ*PSF is the standard deviation of the image PSF approximated by a Gaussian spot; and *σ*_Cam_ is the localization uncertainty.

We estimated the camera-based localization uncertainty for a pixel size of 100 nm and a diffraction-limited PSF at 670 nm wavelength. Fitting the simulated PSF with a 2D Gaussian profile yielded *σ*_PSF_ = 116 nm, whereas measuring the FWHM led to *σ*_PSF_ = 118 nm. Hence, we used *σ*_PSF_ = 117 nm. For the total image background *B* we considered 25 pixels: *B* = 25*b*. The total signal *S* is given by the peak signal *S* and the width of the 2D Gaussian distribution: 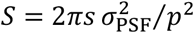. Hence, *S* was defined by the signal fraction of the detected photons *N* = *S* + *B*:

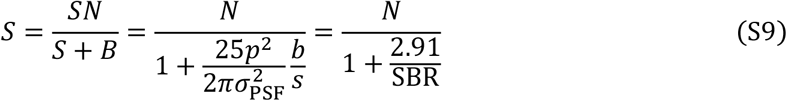

With the chosen imaging parameters, equation (S8) evaluates to

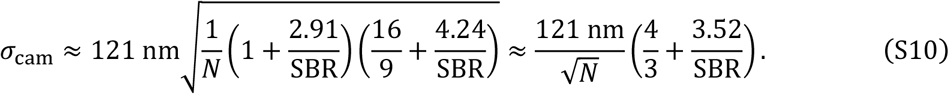

The localization accuracy is further affected by any differences in pixel responses, which is particularly critical when seeking an uncertainty of 10 nm or less with (scientific) CMOS camera images.

#### Comparison of MINSTED and camera-based localization

**Suppl. Fig. S1** illustrates the simulated localization uncertainty for different peak SBRs and the following settings: scan radius *R* = *d*/2 from 103 nm initially down to 13 nm; a centre update step *α* = 15 % of the scan radius; a reduction factor *γ* = 0.97; *ω* = 2*π* × 125 kHz circling frequency; and *ɛ* = 30 kcps peak emission rate, corresponding to about 15–20 kcps average detection rate including background. The localizations were terminated after 10000 photon detections, or earlier when less than *n*_OFF_ = 10 to 15 photons were detected in a *τ*_OFF_ = 3 ms interval. For each setting, we simulated 500 localizations. Less than 3% of the localizations terminated early. If the estimated fluorophore position 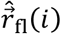 was further off than 5 times the median error of all simulated localizations, we deemed the localization as failure and excluded it for the remaining *i* … *N* photon detections. A fraction of up to 1/SBR localizations failed during the homing-in. An uncertainty of 1 nm along the *x* and *y* directions was obtained with 300 to 800 photon detections.

**Suppl. Fig. S1** also shows the camera-based localization uncertainty without background and for the lowest peak SBR used in the simulations. Dotted lines extend these estimates for low photon numbers that are usually discarded. After a transitory phase, the scanning-based localization massively benefits from the zooming-in with an ever sharper E-PSF. Localization uncertainties of 3 nm or less can be achieved by camera-based localizations but require 30 to 70 times the number of detected photons than MINSTED.

If the fluorophore supports a higher exposure to STED light during the localization, a larger scan radius with respect to the PSF diameter further squeezes the required photon detections. **Suppl. Fig. S2** shows that MINSTED can reach 1 nm precision with as few as 200 detected photons, which is about 100 times more photon-efficient than camera-based localizations and on par with iterative MINFLUX^8^.

#### Animations

**Suppl. Video V1** illustrates the evolutions of the centre-to-fluorophore distance distributions during the detection of 100 photons for update steps *α* = 10%, 15%, 20% and 25% of the scan radius and for peak SBR of 50, 20, 10, 5 and 2. The diameter of the solid circle and the radius of the dashed circle equal the E-PSF diameter *d*, which is twice the scan radius *R*. Each run starts with a uniform centre-to-fluorophore distance distribution in the dashed circle. If the centre-to-fluorophore distance leaves this region, the fluorophore is considered “lost” because the centre would rarely re-approach the fluorophore in practice. Fluorophores get lost mostly during the first few detections when a step in the wrong direction can be fatal. Once the distribution converges, losses occur only if the background is too high and/or the step too large. Towards the end of each run, the standard deviation *σ*_C_ of the converged distribution is shown in units of the scan radius.

**Suppl. Video V2** animates the localization of a fluorophore. Yellow to red dots mark the most recent centre coordinates and the circular line illustrates the recent donut positions. Photon detections are illustrated by a flash with the shape of the E-PSF at the donut position of the detection event. The scan trajectory is updated immediately upon the detection of a photon. Weighted histograms of the centre positions are shown above and to the right of the image. After homing in on the fluorophore, these histograms converge towards normal distributions, whose centres indicate the fluorophore position and whose standard deviations equal the values shown in **Suppl. Video V1**. The animation is sped up about ten-fold by removing full scan circles without detection event.

## Supplementary figures

**Suppl. Fig. S1.**
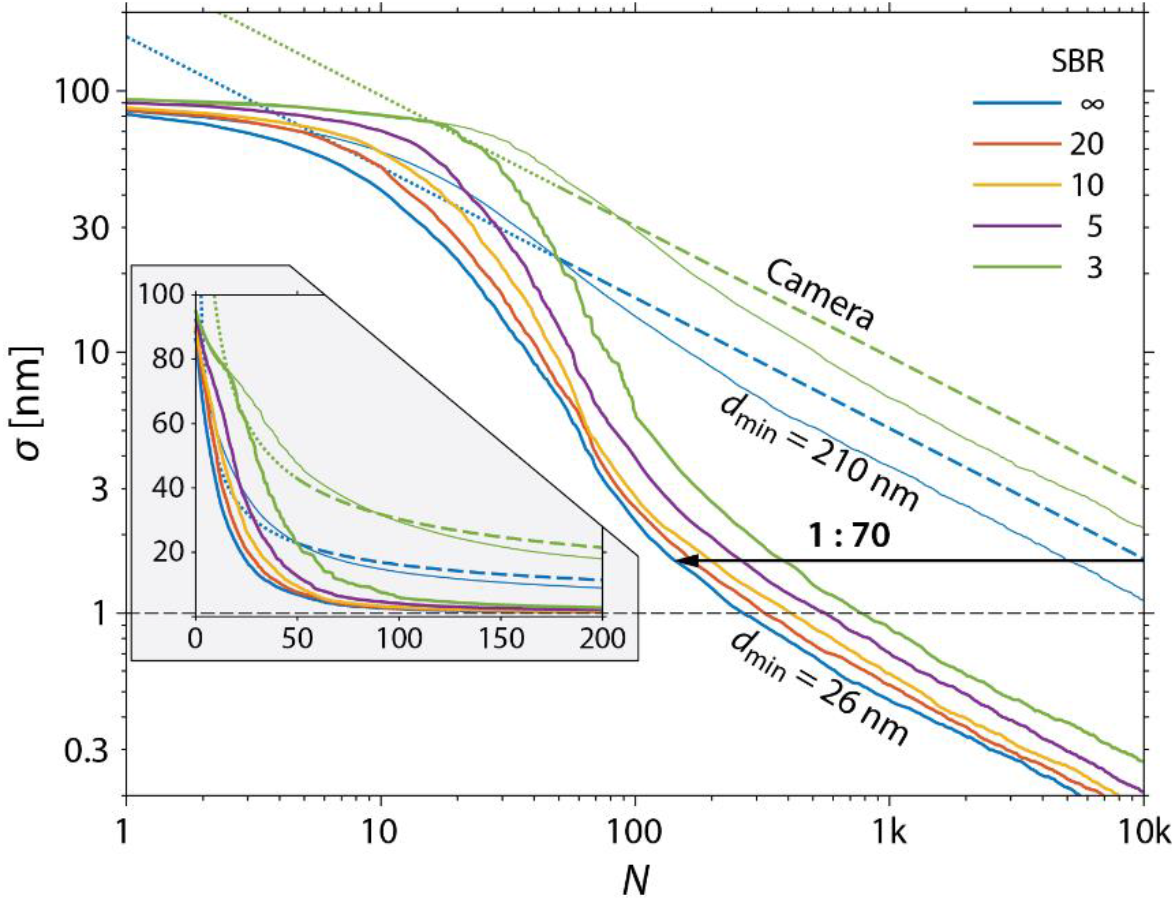
Simulated localization uncertainty versus number of detected photons and SBR. The localizations were performed by zooming-in from *d*_0_ = 210 nm PSF diameter and *R*_0_ = 103 nm scan radius down to *d*_min_ = 26 nm and *R*_min_ = 13 nm. For SRB = ∞ and 3, dashed and dotted lines show the camera-based CRBs and thin lines show the uncertainties with diffraction-limited zooming-in on the fluorophore.

**Suppl. Fig. S2.**
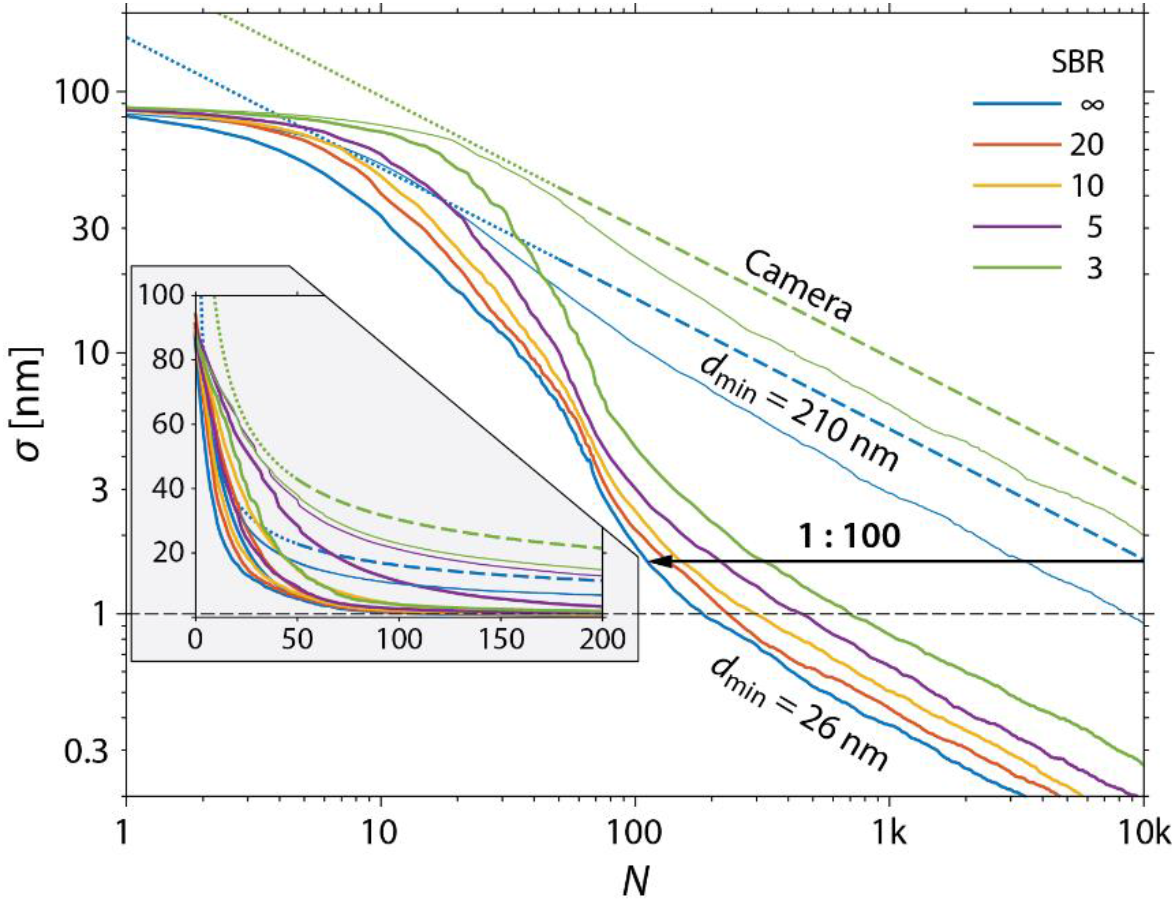
Simulated localization uncertainty versus number of detected photons and SBR. The localizations were performed by zooming-in on the fluorophore from *d*_0_ = 210 nm PSF diameter and *R*_0_ = 130 nm scan radius down to *d*_min_ = 26 nm and *R*_min_ = 17 nm. The inset shows the results in linear scale for small photon numbers. For SRB = ∞ and 3, dashed and dotted lines show the camera-based CRBs and thin lines show the uncertainties with diffraction-limited zooming-in.

**Suppl. Fig. S3.**
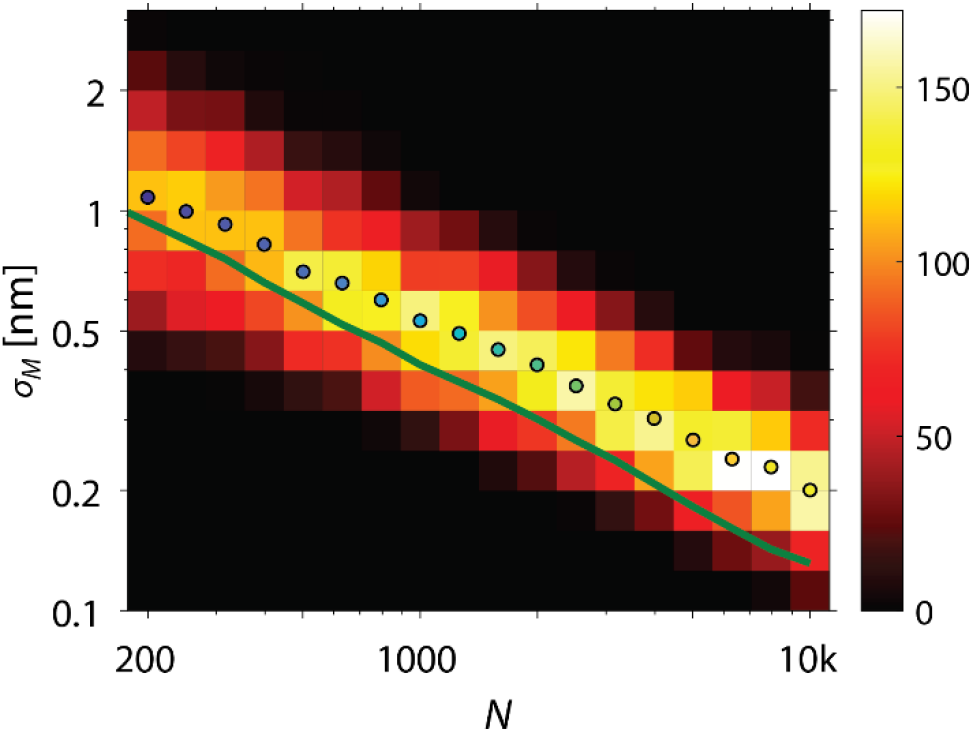
Simulated versus estimated localization precision. The simulated localization uncertainty (solid line) was obtained by Eq. (S2) and the estimated localization precision (histogram; dots: median) by Eq. (S4).

**Suppl. Fig. S4.**
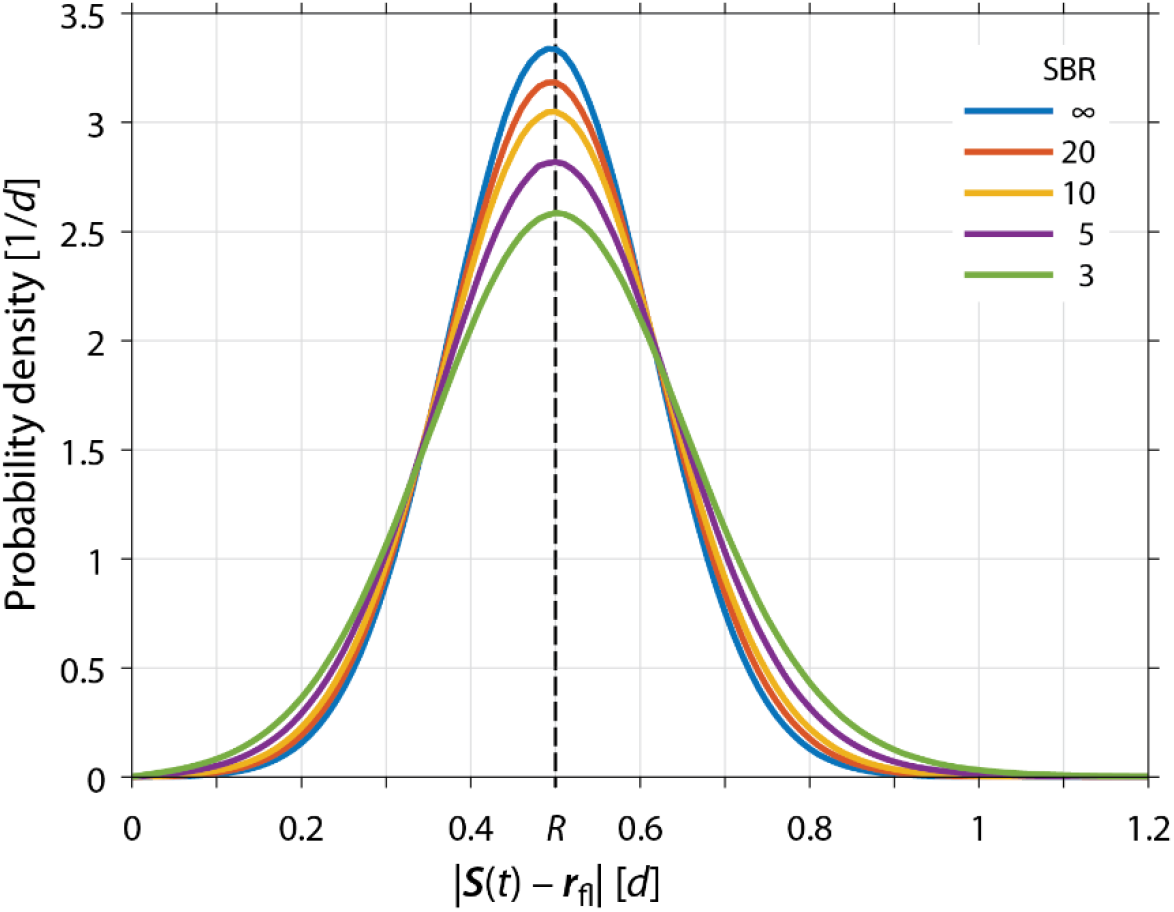
Distribution of the donut-to-fluorophore distances during the simulated localizations of **Suppl. Fig. S1.**

**Suppl. Fig. S5.**
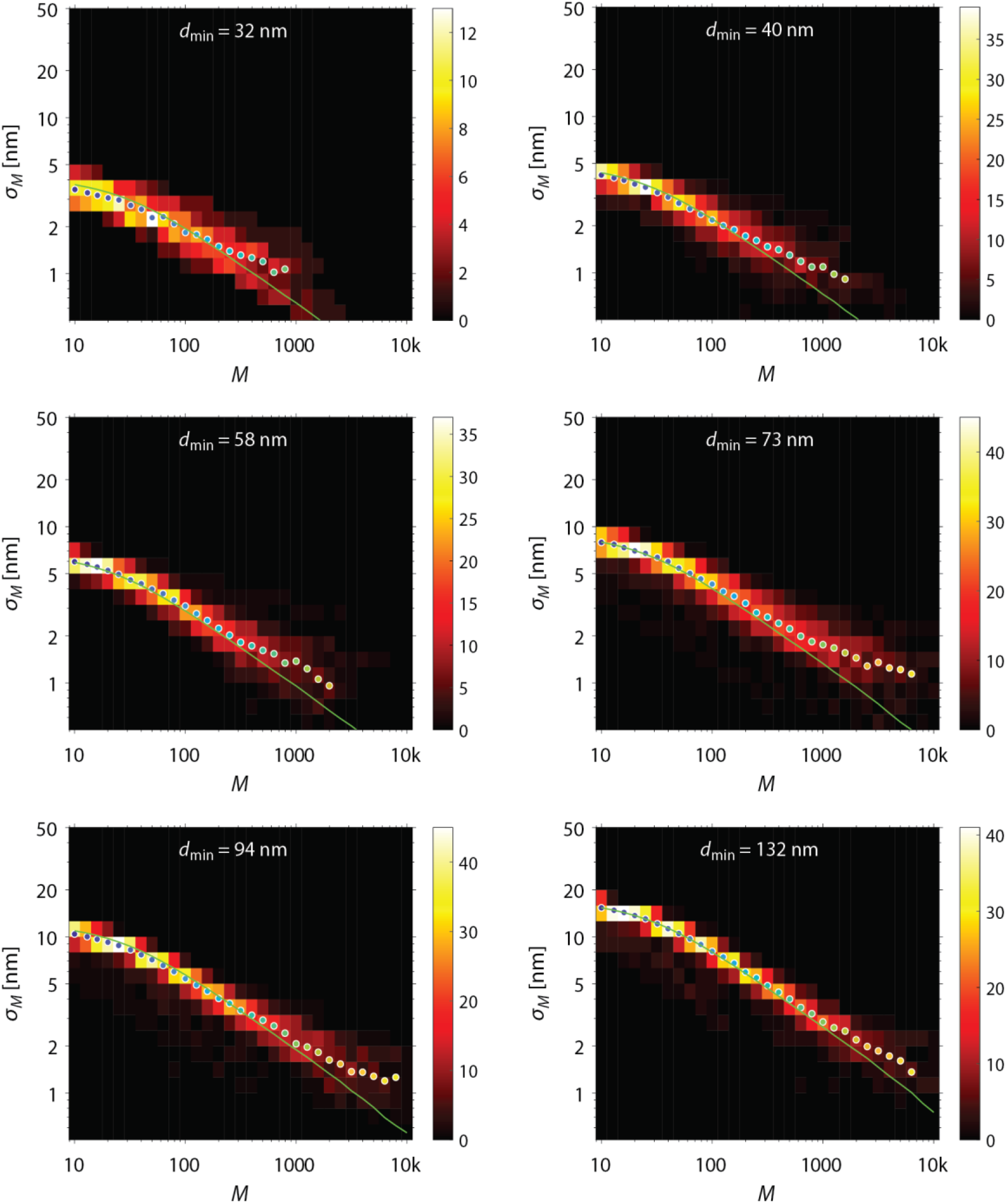
Localization precision measurements versus E-PSF diameter. *d*_min_, see also Fig. 3c. Localization precision histograms of grouped localization traces of single molecules and their median localization precision (dots) compared to simulations (lines).). The simulations assumed an SBR of 10 for *d*_min_ = 32, 94 and 132 nm, an SBR of 20 for *d*_min_ = 40 and 73 nm and infinite for *d*_min_ = 58 nm.

**Suppl. Fig. S6.**
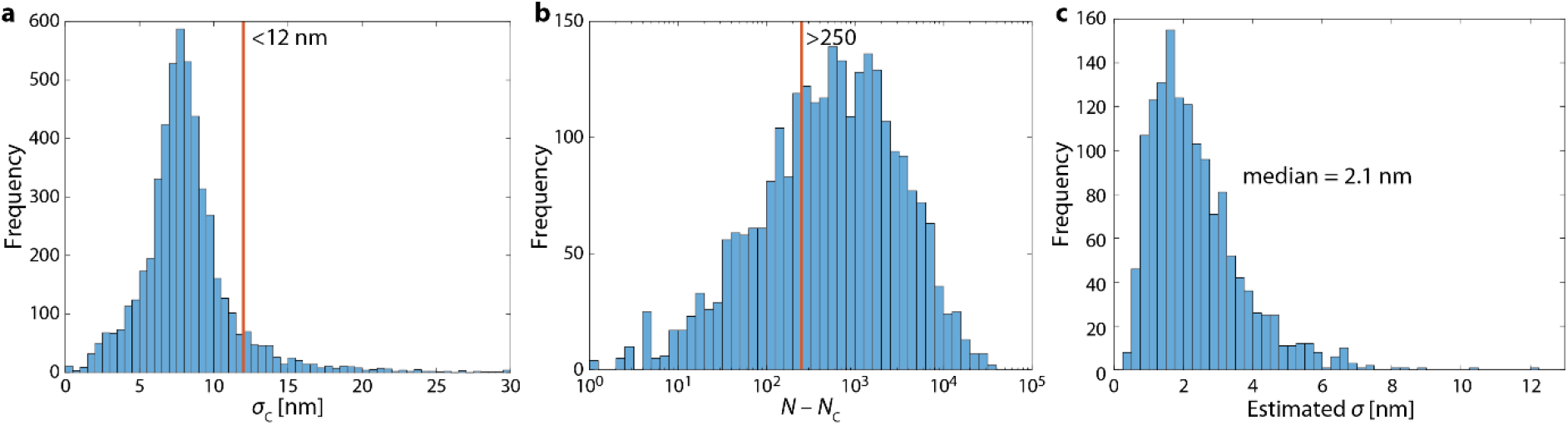
Characteristics of the localizations measured for Fig. 4. **a**, Distribution of the standard deviation of the centre positions. **b**, Distribution of the number of detected photons used for estimating the fluorophore position. **c**, Distribution of the estimated localization precision of the rendered localizations.

## Supplementary videos

**Suppl. Video V1.**
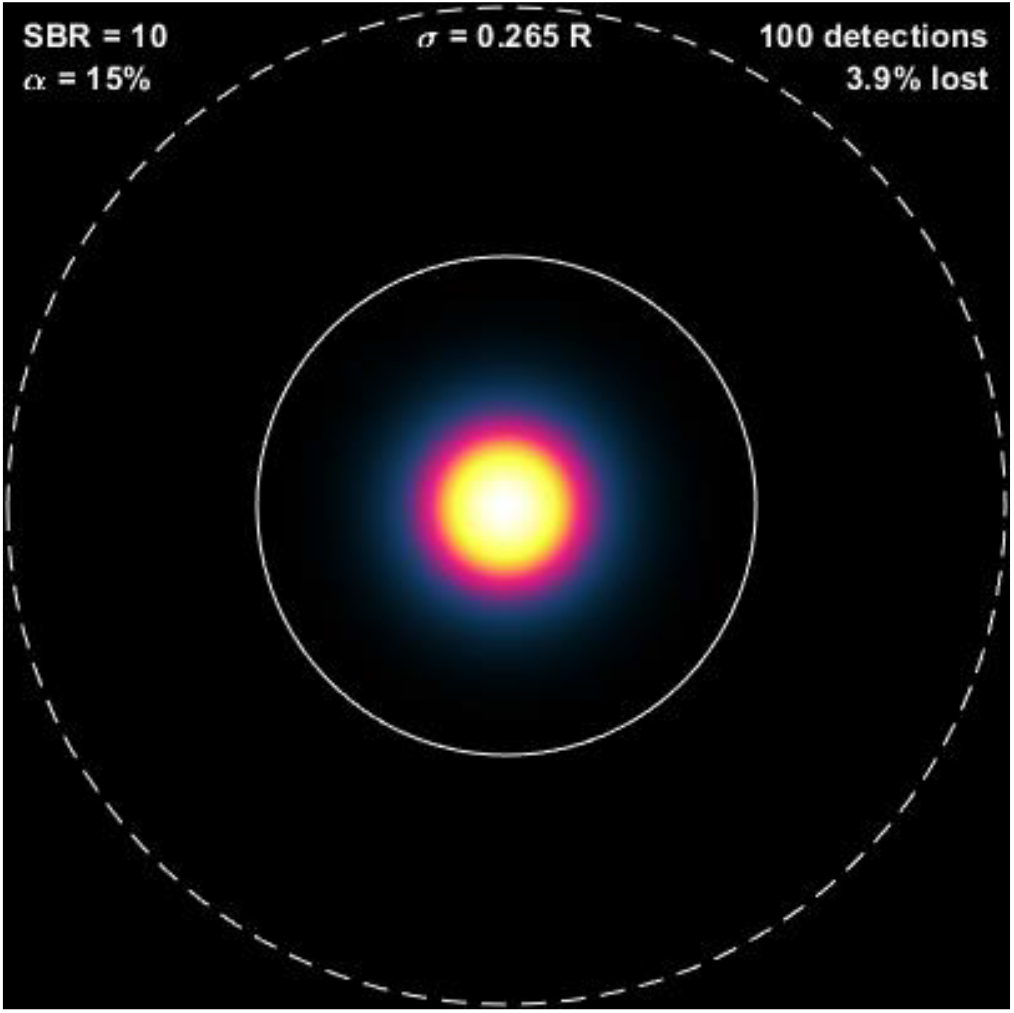
Distributions of the centre-to-fluorophore distances during the detection of 100 photons while circling with constant scan radius and constant E-PSF. The solid circle illustrates the scan trajectory and the E-PSF diameter.

**Suppl. Video V2.**
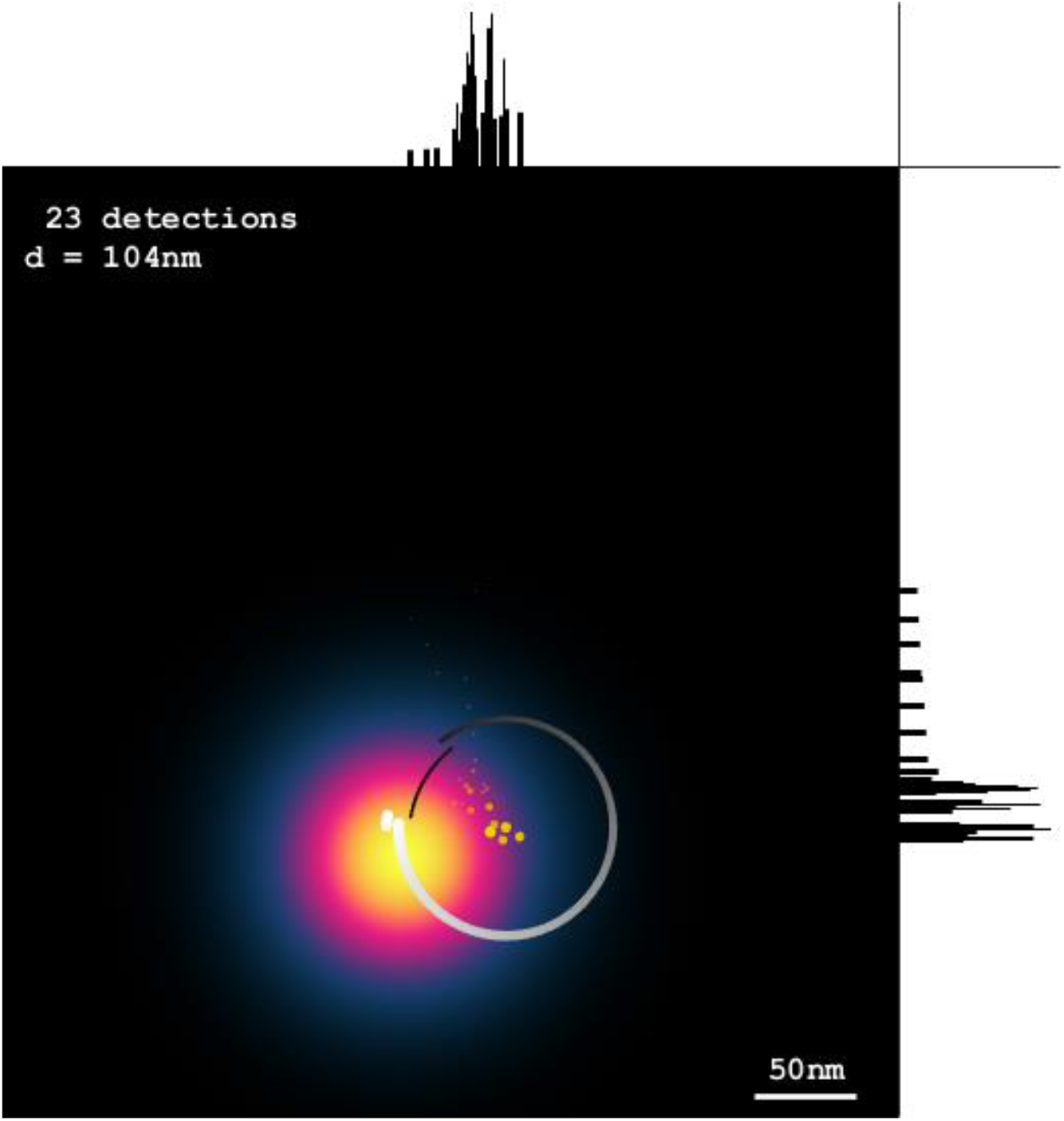
Animation of the localization of a single fluorophore with typical settings: scan radius *R* = d/2, update steps *α* = 15% and *γ* = 0.97.

## References

1. Hell, S. W. & Wichmann, J. Breaking the diffraction resolution limit by stimulated emission: stimulated-emission-depletion fluorescence microscopy. Opt. Lett. 19, 780–782 (1994).

2. Klar, T. A. et al. Fluorescence microscopy with diffraction resolution barrier broken by stimulated emission. Proc. Natl. Acad. Sci. USA 97, 8206–8210 (2000).

3. Westphal, V. & Hell, S. W. Nanoscale Resolution in the Focal Plane of an Optical Microscope. Phys. Rev. Lett. 94, 143903 (2005).

4. Hell, S. W. Far-Field Optical Nanoscopy. Science 316, 1153–1158 (2007).

5. Danzl, J. G. et al. Coordinate-targeted fluorescence nanoscopy with multiple off states. Nature Photon. 10, 122–128 (2016).

6. Betzig, E. et al. Imaging Intracellular Fluorescent Proteins at Nanometer Resolution. Science 313, 1642–1645 (2006).

7. Rust, M. J., Bates, M. & Zhuang, X. Sub-diffraction-limit imaging by stochastic optical reconstruction microscopy (STORM). Nature Methods 3, 793–796 (2006).

8. Balzarotti, F. et al. Nanometer resolution imaging and tracking of fluorescent molecules with minimal photon fluxes. Science 355, 606–612 (2017).

9. Levi, V., Ruan, Q., Kis-Petikova, K. & Gratton, E. Scanning FCS, a novel method for three-dimensional particle tracking. Biochem. Soc. Transact. 31, 997–1000 (2003).

10. Göttfert, F. et al. Strong signal increase in STED fluorescence microscopy by imaging regions of subdiffraction extent. Proc. Natl. Acad. Sci. USA 114, 2125–2130 (2017).

11. Heine, J. et al. Adaptive-illumination STED nanoscopy. Proc. Natl. Acad. Sci. USA 114, 9797–9802 (2017).

12. Kasper, R. et al. Fluorophores: Single-Molecule STED Microscopy with Photostable Organic Fluorophores. Small 6, 1379–1384 (2010).

13. Weber, M. et al. Photoactivatable Fluorophore for STED Microscopy and Bioconjugation Technique for Hydrophobic Labels. Chem. Eur. J. (2020). DOI: 10.1002/chem.202004645

14. Pfanner, N. et al. Uniform nomenclature for the mitochondrial contact site and cristae organizing system. J. Cell. Biol. 204, 1083–1086 (2014).

15. Jans, D. C. et al. STED super-resolution microscopy reveals an array of MINOS clusters along human mitochondria. Proc. Natl. Acad. Sci. USA 110, 8936–8941 (2013).

16. Saphire E. O. et al. Crystal Structure of a Neutralizing Human IgG Against HIV-1: A Template for Vaccine Design. Science 293, 1155–1159 (2001).

17. Pape, J. K. et al. Multicolor 3D MINFLUX nanoscopy of mitochondrial MICOS proteins. Proc. Natl. Acad. Sci. USA 117, 20607–20614 (2020).

18. Gwosch, K. C. et al. MINFLUX nanoscopy delivers 3D multicolor nanometer resolution in cells. Nature Methods 17, 217–224 (2020).

19. Rittweger, E. et al. STED microscopy reveals crystal colour centres with nanometric resolution. Nature Photon. 3, 144–147 (2009).

20. Puthukodan, S., Murtezi, E., Jacak, J. & Klar, T. A. Localization STED (LocSTED) microscopy with 15 nm resolution. Nanophoton. 9, 783–792 (2020).

## References

21. Vogelsang, J. et al. A Reducing and Oxidizing System Minimizes Photobleaching and Blinking of Fluorescent Dyes. Angew. Chem. Int. Ed. 47, 5465–5469 (2008).

## Supplementary references

22. Thompson, R. E., Larson, D. R. & Webb, W.W. Precise nanometer localization analysis for individual fluorescent probes. Biophy. J. 82, 2775–2783 (2002).

23. Mortensen, K.I. et al. Optimized localization analysis for single-molecule tracking and super-resolution microscopy. Nature Methods 7, 377–381 (2010).

